# Non-Newtonian Blood Rheology Significantly Alters Hemodynamic Predictions During Cardiac Looping: A Computational Study

**DOI:** 10.64898/2026.05.15.725470

**Authors:** C. Matthew, Erica C. Kemmerling, Lauren D Black

**Author notes:** **Corresponding Author:** Lauren D. Black III, Department of Biomedical Engineering, Tufts University, 4 Colby Street, Medford, MA 02155. Authors contributed equally to this work (Erica C. Kemmerling, Lauren D Black).

## Abstract

Hemodynamic forces play a key role in early cardiac morphogenesis, yet many computational studies assume Newtonian blood behavior. Here, we evaluate the impact of non⍰Newtonian shear⍰thinning rheology on flow patterns, pressure distributions, and wall shear stress (WSS) during cardiac looping using idealized three⍰dimensional models of the embryonic heart tube. Five geometries representing progressive looping stages, from a linear tube to an S⍰shaped configuration with ventricular ballooning, were analyzed under pulsatile flow using both Newtonian and power⍰law viscosity models. Across all stages, Reynolds numbers (Re ≈ 1–7) and Womersley numbers (Wo ≈ 0.3) indicated laminar, quasi⍰steady flow consistent with embryonic conditions. Incorporating shear⍰thinning rheology produced substantial deviations from Newtonian predictions, with peak systolic WSS differing by up to ∼40% and pressure drops by up to ∼20%. These effects were most pronounced in regions of increased curvature and geometric complexity. These findings demonstrate that non⍰Newtonian rheology significantly influences predicted hemodynamic environments during cardiac looping and should be incorporated into computational models aimed at understanding mechanobiological regulation of early heart development.

## INTRODUCTION

The heart is the first functioning organ to form in embryonic vertebrates—its first contractions begin around day 21 of embryonic development in humans [1–4]. The heart begins as a linear tube, consisting of a single layer of endothelial cells surrounding the lumen and a layer of primitive cardiac extracellular matrix, called cardiac jelly, separating the endothelium from an outer layer of beating myocytes. Over several days, the linear heart tube undergoes a process called “cardiac looping” to transform into its mature, four-chambered shape. While genetic and biological cues also play a role in proper heart formation and function, it has long been speculated that hemodynamics modulate cell signaling and tissue function via physical cues such as wall shear stress. In many cases, altered blood flow has been correlated with pathological heart formation, resulting in defects such as hypoplastic left heart syndrome [3, 5, 6]. While many studies have focused on hemodynamics in the developing ventricle and chambered hearts [7–10], fewer studies have examined blood flow in the primitive embryonic heart tube. An accurate understanding of hemodynamics throughout cardiac looping could lead to a more complete understanding of cardiac development and could help elucidate cardiac abnormalities and pathologies that occur during embryonic development.

There are many physical forces present during cardiac looping, and the direct mechanical methods that drive mechanical looping remain poorly understood [11]. Blood flow begins around the third gestational week in humans while the heart is a linear tube without valves or chambers [3]. Blood flow exerts two stresses—wall shear stress and hydrostatic pressure—on the early heart walls [5]. Cardiac cells can sense and respond to these stresses, and deviations from normal physiological stresses can result in serious pathologies [6, 12, 13]. Thus, an understanding of the magnitudes of these stresses and how they change throughout embryonic cardiac development could help elucidate the causes of pathologies and offer insight into corrective procedures.

It has been well established that blood flow is critical for healthy cardiac development; however, it is difficult for *in vivo* experiments to capture the fluid mechanics present during heart development in humans. Computational fluid dynamics (CFD) simulations have emerged as a promising tool evaluate cardiac flow features such as wall shear stresses and pressure distributions. In recent years, CFD has been a tool to investigate blood flow during congenital heart disease [14, 15], as well as for studying the underlying fluid mechanics present during cardiac development [16]. Most CFD models which investigate embryonic heart looping assume a Newtonian model of blood, due to the low hematocrit of embryonic blood. In adult vertebrates, hematocrits are usually greater than 40% [17], and non-Newtonian behavior is exacerbated due to increased interactions between deformable red blood cells. Embryonic blood has a hematocrit that ranges between 20%-30% [18], lower than adult blood, but still non-Newtonian in behavior. Previous models have shown that using a Newtonian model in a physiologically-motivated geometry can result in large errors in flow solutions [19]. For example, in straight tubes, blunted velocity profiles and higher pressure gradients are observed in shear-thinning simulations compared to Newtonian simulations. Additionally, shear thinning flows in curved tubes have weaker secondary flows and less axial vorticity than Newtonian flows [19]. We believe that it is necessary to incorporate a non-Newtonian model of blood and pulsatile flow to better simulate the fluid dynamics present during embryonic heart looping. In this study, we present a CFD model that interrogates differences in hemodynamics during different stages of embryonic heart tube looping, specifically investigating differences in wall shear stress and pressure. We present data on the wall shear stresses and pressure distributions in non-Newtonian and Newtonian models of blood flow during embryonic heart looping.

## METHODS

### Heart tube stage geometries

Most CFD models use optical techniques, such as magnetic resonance imaging, cardio echography, or line scan confocal microscopy, to generate models of embryonic ventricles and animal models to obtain similar geometries [16]. While these optical techniques make it possible to generate complex models from late-stage embryonic heart ventricles, to the authors knowledge, no techniques exist that have the resolution to accurately reconstruct the small sized early human heart tube stages *in utero*. Thus, our model uses idealized geometries, representative of dimensions of the developing embryonic heart tube, to study the hemodynamics present within early stages of cardiac looping. Idealized geometries have been used before to study embryonic heart tube development. For example, Taber et al. modeled the transition from oscillatory blood flow to pulsatile blood flow within an idealized two dimensional linear heart tube model with symmetric cardiac cushions [20].

We used idealized geometries representing five stages of cardiac looping, starting from a linear tube, transitioning through a C-shaped tube, and finally forming an S-shaped tube (Figure 1, Table 1). Idealized geometries were intentionally selected to enable systematic, stage-by-stage comparison of hemodynamic changes arising from looping curvature, tube elongation, and ventricular ballooning, independent of inter-individual geometric variability inherent to image-based reconstructions. This approach is particularly important for early human heart tube stages, for which current in utero imaging techniques lack sufficient spatial and temporal resolution. Maximum lumen diameters and lengths (Table 1) were chosen to be representative of reported dimensions of the developing human embryonic heart tube [7, 20], with lumen diameters increasing across stages to capture ventricular ballooning during looping. The largest lumen diameter (1 mm) in the Stage 5 model is comparable to reported chamber dimensions in early human embryonic ventricles [7]. Similar idealized approaches have been widely used to interrogate mechanistic drivers of embryonic cardiac development [20], enabling controlled evaluation of specific geometric and rheological effects that would be difficult to isolate in anatomically reconstructed models.

**Figure 1:**
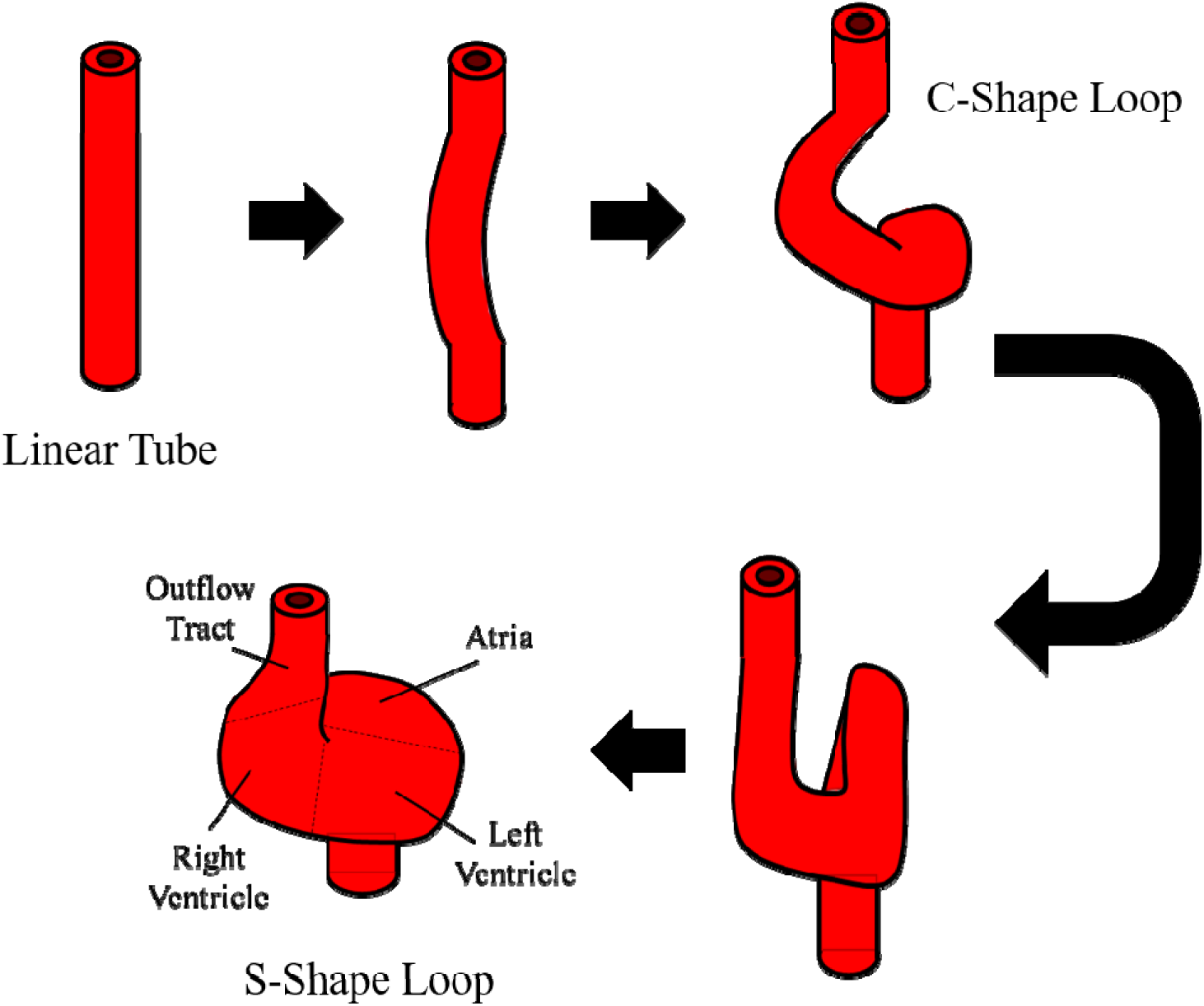
Schematic of the transition from the linear heart tube into, C-stage, and Sstage heart tube geometries. In addition to looping, the heart tube balloons and lengthens with the proliferating, stretching, and migrating cells.

**Table 1:**
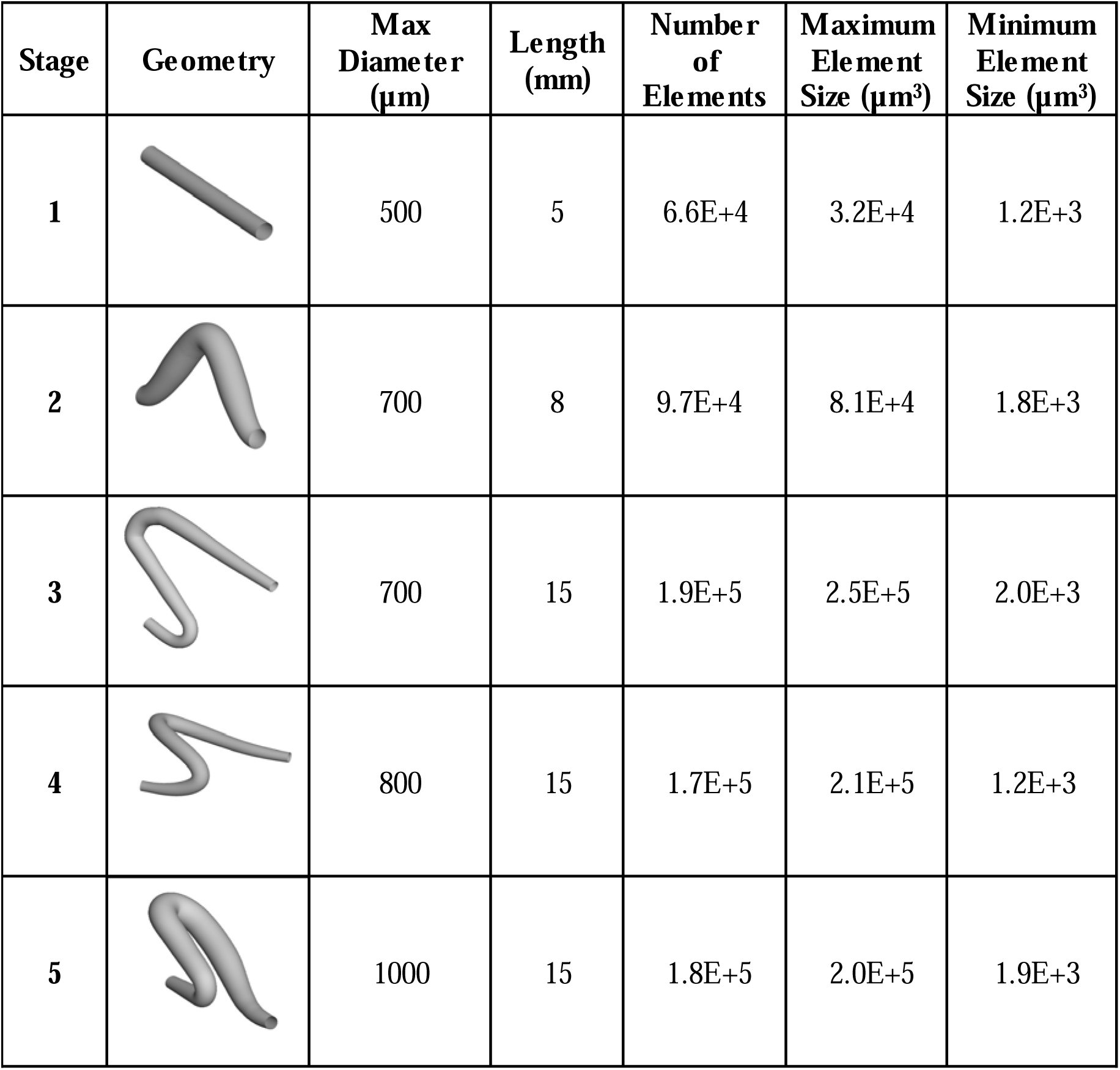
Summary of geometries and meshes.

| Stage | Geometry | Max Diameter ($\mu\text{m}$) | Length (mm) | Number of Elements | Maximum Element Size ($\mu\text{m}^3$) | Minimum Element Size ($\mu\text{m}^3$) |
| --- | --- | --- | --- | --- | --- | --- |
| 1 |  | 500 | 5 | 6.6E+4 | 3.2E+4 | 1.2E+3 |
| 2 |  | 700 | 8 | 9.7E+4 | 8.1E+4 | 1.8E+3 |
| 3 |  | 700 | 15 | 1.9E+5 | 2.5E+5 | 2.0E+3 |
| 4 |  | 800 | 15 | 1.7E+5 | 2.1E+5 | 1.2E+3 |
| 5 |  | 1000 | 15 | 1.8E+5 | 2.0E+5 | 1.9E+3 |

### Meshing

Table 1 summarizes the meshes and geometries for the five stages. Geometries were imported into ANSYS v19.2 workbench and meshes were generated for each stage using the ANSYS meshing software. Table 1 summarizes the number of elements for each geometry. Structured meshes featuring hexahedral elements with inflation near the lumen walls were used for all geometries. First, a growth layer five cells deep with 50 cells in the circumferential direction was applied to the face of the inlet boundary. The thinnest cell was at the edge and had a thickness of 3 µm. Inner cells had a growth rate of 1.2 in the radial direction. Mesh volumes were generated by sweeping the face grid along the outer radius of the geometries. A grid convergence study was performed to determine the minimum number of elements needed for each solution to have less than 1% error.

### Blood material properties

Blood is a complex fluid that contains deformable cells. Despite this, many blood simulations have approximated blood as Newtonian. Rather, blood is non-Newtonian, and exhibits both shear-thinning and viscoelastic behavior due to interactions between red blood cells. Different methods have been used to model the non-Newtonian properties of blood. One common method is to model blood as a two-phase material featuring a Newtonian boundary layer without cells, and a core region rich with red blood cells [21]. Red blood cells naturally accumulate in the center of blood vessels, a phenomenon called the Fahraeus-Lindqvist effect. Two-phase models accurately capture this effect [21]; however, piecewise defined viscosity distributions are prone to discontinuous derivatives at the phase boundaries [19], and can lead to unstable solutions.

Another common technique for modeling blood viscosity is continuous viscosity functions [19, 22]. These functions assign a viscosity to every point in the domain based on local shear rate, position, or time history. Continuous viscosity models have a relatively low computational cost, and can model both shear and viscoelasticity [19, 23, 24]. In our model, the viscosity of blood was modeled as a function of local shear rate. A power function (Eq. 1) was used to fit data of shear rate dependence of blood viscosity for blood with a hematocrit of 28% from Brooks et al. [18].

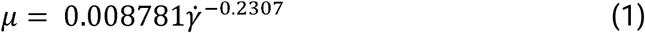

Here, *µ* is the dynamic viscosity in *Pa·s* and γ̇ is the local shear rate in 1/s. For shear rates less than 0.1 s^−1^, the viscosity was set to be a constant 0.0149 Pa·s. A user defined function prescribed a viscosity value for each grid element within our geometries. Briefly, during each iteration, the local shear rate was calculated within each cell, and the viscosity of the fluid within each element was adjusted based on our power fit viscosity model.

Blood also has viscoelastic characteristics. The Weissenberg number (Wi) is a dimensionless number used to characterize viscoelastic flows (Eq. 2) and can be interpreted as the ratio of elastic to viscous forces [25].

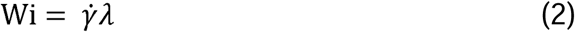

Here, *γ̇* is defined as the shear rate and *λ* is the relaxation time [25]. Relaxation time is defined as the amount of time a strained viscoelastic material takes to return to its initial equilibrium. Extensive rheology experiments conducted by Thurston *et al*. investigated Wi and relaxation times for blood in oscillatory flow in different flow geometries. For 22% hematocrit blood in a 0.10 cm diameter tube with an oscillatory frequency of 2 Hz, the maximum Wi was observed to be 0.1 at a shear rate of 12.7 s^−1^. Wi leveled off to 0.07 at shear rates greater than 100 s^−1^ [17]. Based on the data from Thurston *et al*. [17] and and estimated shear rates within the developing embryonic heart tube, we expect Weissenberg numbers (Wi) to vary between 0 and approximately 0.1. This indicates that viscous forces dominate elastic effects under the flow conditions studied. Neglecting elastic stress terms within the governing equations therefore introduces a maximum expected error on the order of 10%. Notably, this error is substantially smaller than the differences observed between Newtonian and non Newtonian shear thinning models, which exceeded 40% for wall shear stress predictions at peak systolic flow. Consequently, a shear rate dependent viscosity model captures the dominant non Newtonian behavior of embryonic blood while maintaining numerical stability, computational efficiency, and interpretability of hemodynamic outcomes.

### Governing equations and boundary conditions

The Navier-Stokes equations and continuity equations were solved using ANSYS Fluent. Boundary conditions and initial conditions were applied to our models for each case. For all cases, a no-slip boundary condition was used at the walls. The inlet velocity profile was specified as paraboloid-shaped with a transient flow rate at the inlet for both Newtonian and non-Newtonian models. A fully developed paraboloid inlet velocity profile was assumed to provide controlled and reproducible inflow conditions for systematic comparison across looping stages and viscosity models. Although embryonic inflow is generated by upstream peristaltic motion in a short, valveless heart tube, prior experimental and computational studies indicate that under the low to moderate Womersley numbers characteristic of early embryonic cardiac flow, velocity profiles rapidly approach quasi parabolic shapes over short axial distances [20, 26]. Because all simulations applied identical inflow assumptions, relative comparisons of wall shear stress and pressure distributions between Newtonian and non-Newtonian cases remain robust, even if absolute inflow profile details vary in vivo. A mass flow rate vs. time waveform was obtained from previously published data by Taber et al. [20] and used as the flow rate waveform for our models (Figure 2 A). Blood flow through the heart tube increases throughout cardiac looping and embryonic development (Figure 2 B), and we accounted for this increase by scaling the mass flow rate waveform with published data of cardiac output data vs. gestational week [27] (Figure 2 C). Once the average mass flow rate of one pulse was determined for each stage, the average velocity vs. time was determined by dividing the flow rate by the cross-sectional area of the corresponding model. For all models, the derivative of pressure at the outlet was 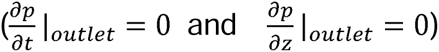, and all pressures within the fluid domain are reported relative to the constant outlet pressure [1]. While cardiac output data were extrapolated to approximate early gestational stages, all simulations preserve consistent waveform shape and relative scaling across looping stages, ensuring that comparative analyses between viscosity models are internally consistent even if absolute flow magnitudes vary in vivo.

**Figure 2:**
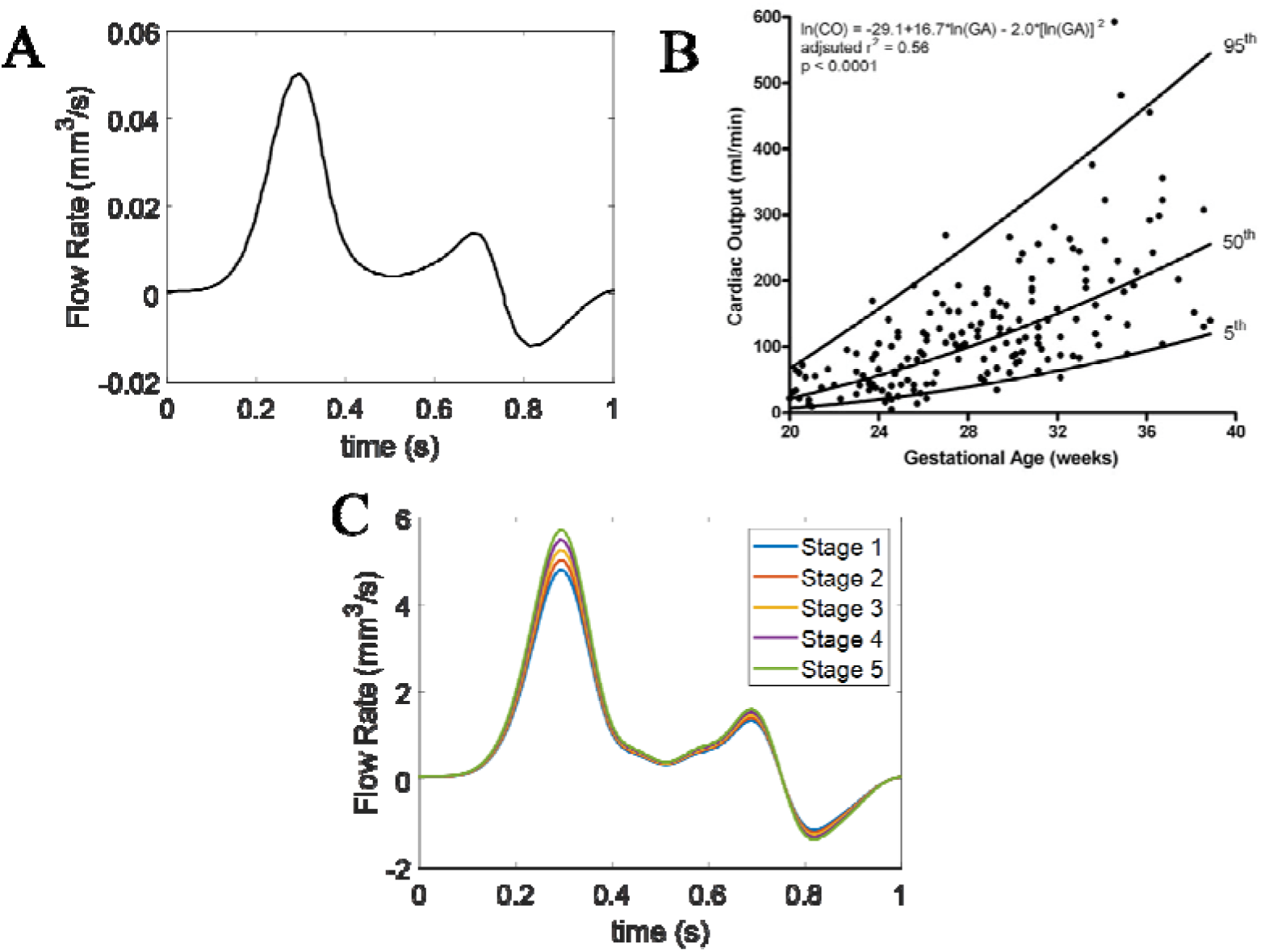
(A) Pulsatile flow rate computed from the outlet of a computational model of flow through a model of the chick embryonic linear heart tube (Taber 2007). (B) Data and fit of cardiac output vs. gestational age for healthy human patients between 20 and 40 weeks (Hamill 2011). (C) Flow rate waveforms used by our models, matching the shape of the embryonic chick outflow waveform and the average cardiac output data extrapolated to three weeks.

### Numerical methods

All simulations were performed in the commercial software Fluent using a second order upwind solver. For our models, we set the time step to be 0.002 s. All simulations were run for 500 time steps. Scaled residuals are defined the imbalance between the left and right sides of the governing equations normalized by the largest error calculated during the first five iterations of each time step. For each time step, simulations were considered converged when scaled mass continuity residuals had converged to 1E-5% error.

### Post-processing

Data was saved as ANSYS data and case files every ten time-steps for each case. Once simulations finished running, data was loaded into Tecplot 360 EX (version 2019 R1). Contour plots of wall-shear stress and hydrostatic pressure were generated for all geometries. Centerline coordinates of each geometry were extracted and used to create slices along inlet, outlet, and bends of each geometry. Contour plots of velocity and viscosity were generated for each slice.

## Results

### Validation of viscosity and velocity implementation

Contour plots of viscosity values were generated for both Newtonian and non-Newtonian cases of the mid cross-sectional plane of our Stage 1 geometry at peak systolic flow. Viscosity remained a constant 0.004 Pa·s for the Newtonian case, while the non-Newtonian case generated low viscosity values near the walls, and high viscosities near the center (Figure 3A-B). Vertical lines were drawn along the diameters of the mid cross-sectional planes for both cases. Velocity data were probed along the lines to generate velocity profiles. As expected, a parabolic shape is observed for the Newtonian model of flow Stage 1 geometry, while the non-Newtonian model featured a blunter and wider velocity profile (Figure 3 C). This is expected for the non-Newtonian case: viscosities are low at high shear rates near the wall and high at low shear rates near the center of the vessel.

**Figure 3:**
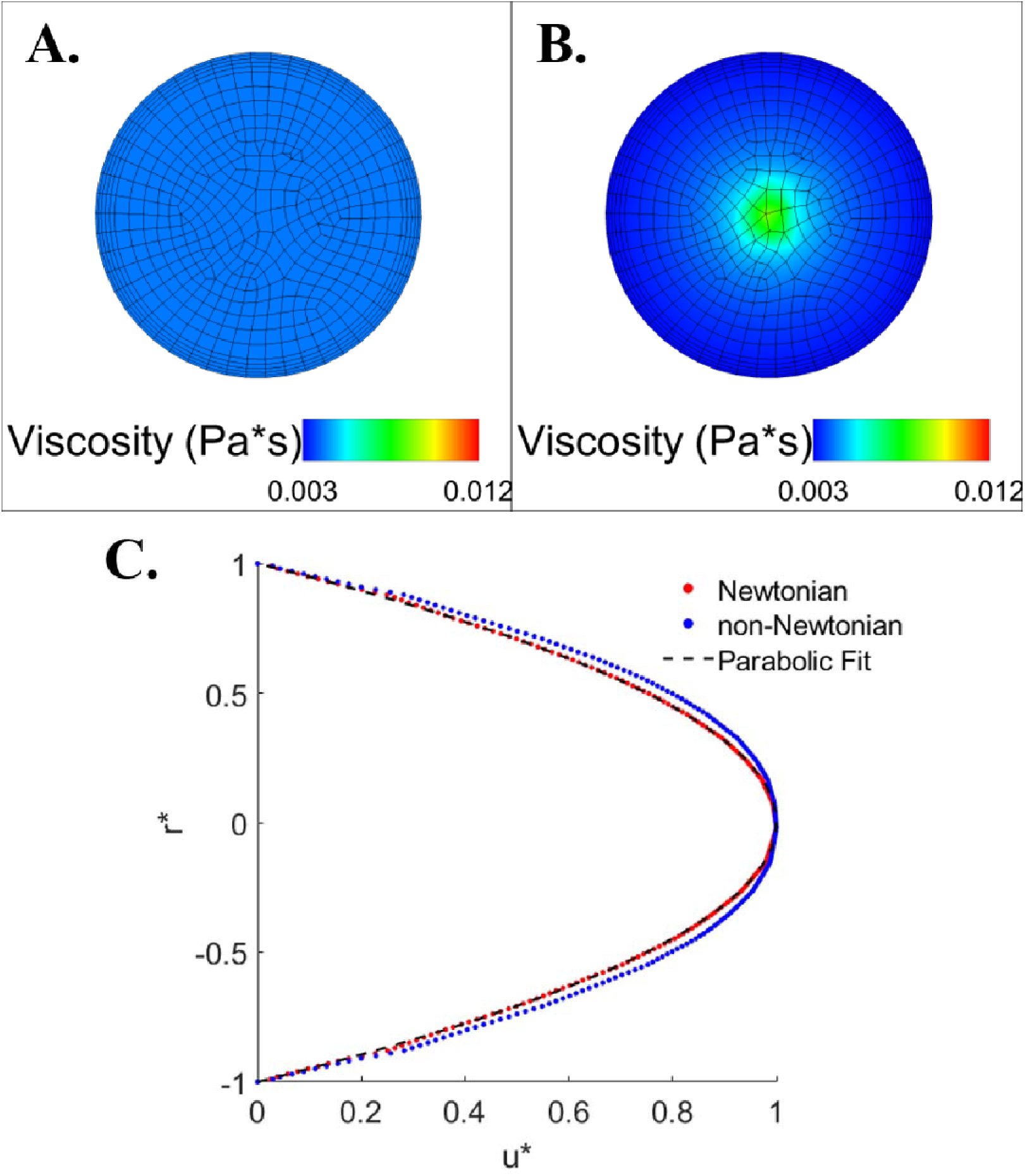
(A) Viscosity value of 0.004 Pa·s at peak diastolic flow through the Stage 1 Newtonian case. (B) Viscosity contours at peak diastolic flow through the Stage 1 non-Newtonian case. Contour plots were generated from data collect at the midway cross-section (*x* = 2.5 mm). (C) Velocity profiles for Newtonian and non-Newtonian Stage 1 cases at peak diastolic flow. *r*\* is the normalized radial position defined *r*/*r_max_* where *r_max_* is the maximum radial postion. *u*\* is the normalized axial velocity defined by *u/u_max_* where *u_max_* is the maximum axial velocity.

### Hydrostatic pressure is overpredicted by Newtonian models

Contour plots of hydrostatic pressures were generated for both Newtonian and non-Newtonian cases at peak systolic flow and peak reverse diastolic flow (Figure 4 and Figure 5). In all models, we observed a monotonic pressure decrease from the inlet to the outlet. This is expected because the pressure gradient must balance viscous forces. Simulations predicted a decrease in pressure between stage 1 and stage 2 geometries possibly due to increased luminal diameters. Pressure drops increased between stage 2 and 3 geometries as expected with an increase in tube length. Finally, pressures dropped between stage 3 to stage 5 geometries where the tube length remained the same and lumen diameters increased. For all cases, Newtonian models of blood predicted higher pressure drops compared to non-Newtonian models. This is expected because lower viscosity values at high shear rates reduce flow resistance and pressures. Maximum pressure values for each stage and viscosity model at peak systolic flow are summarized in table 2.

**Figure 4:**
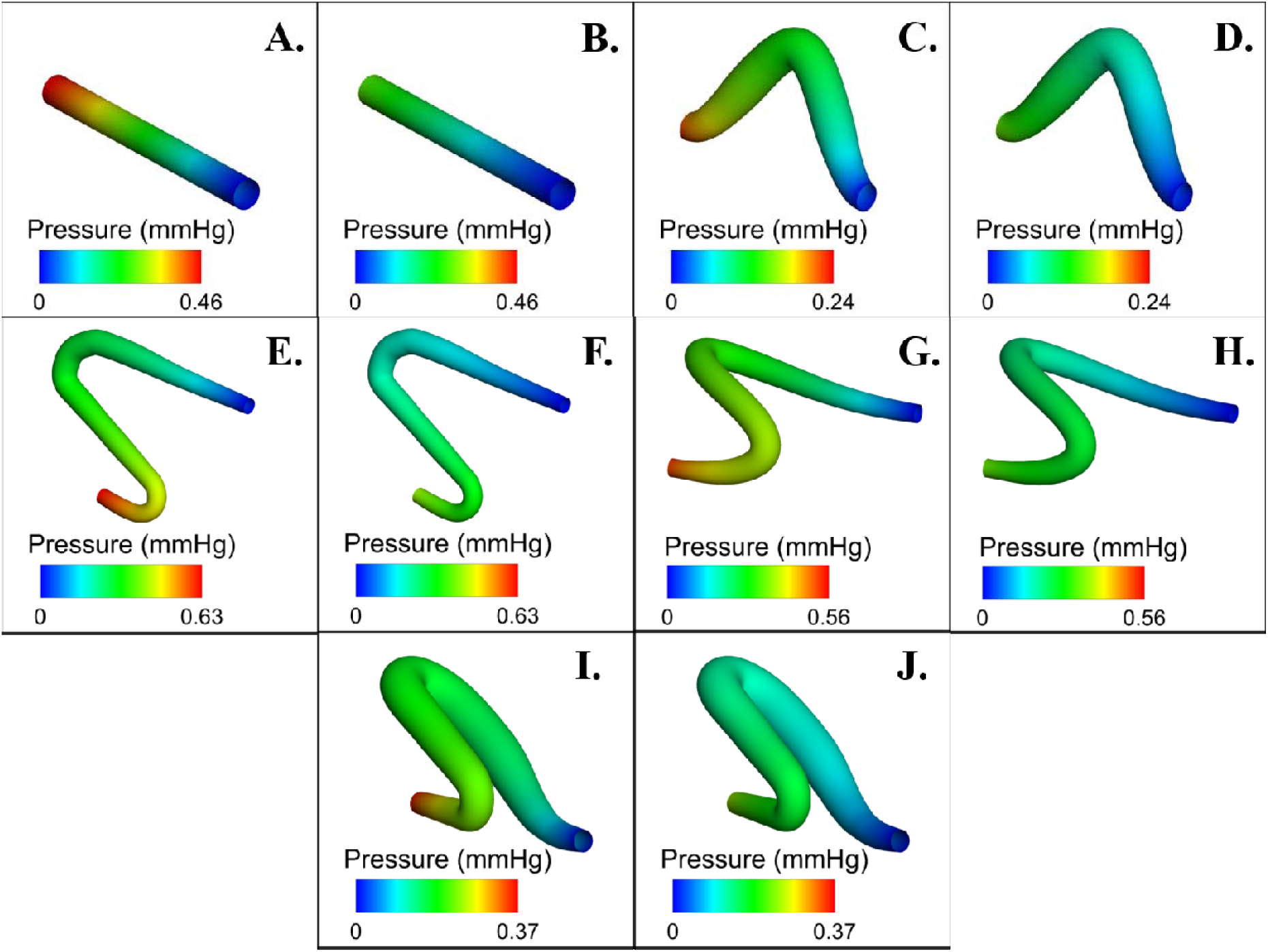
Peak systolic (t = 0.30s) pressure values. Newtonian blood models: (A) Stage 1 (C) Stage 2 (E) Stage 3 (G) Stage 4 (I) Stage 5. Non-Newtonian blood models: (B) Stage 1 (D) Stage 2 (F) Stage 3 (H) Stage 4 (J) Stage 5. For all geometries, the inlet diameters are 0.500 mm.

**Figure 5:**
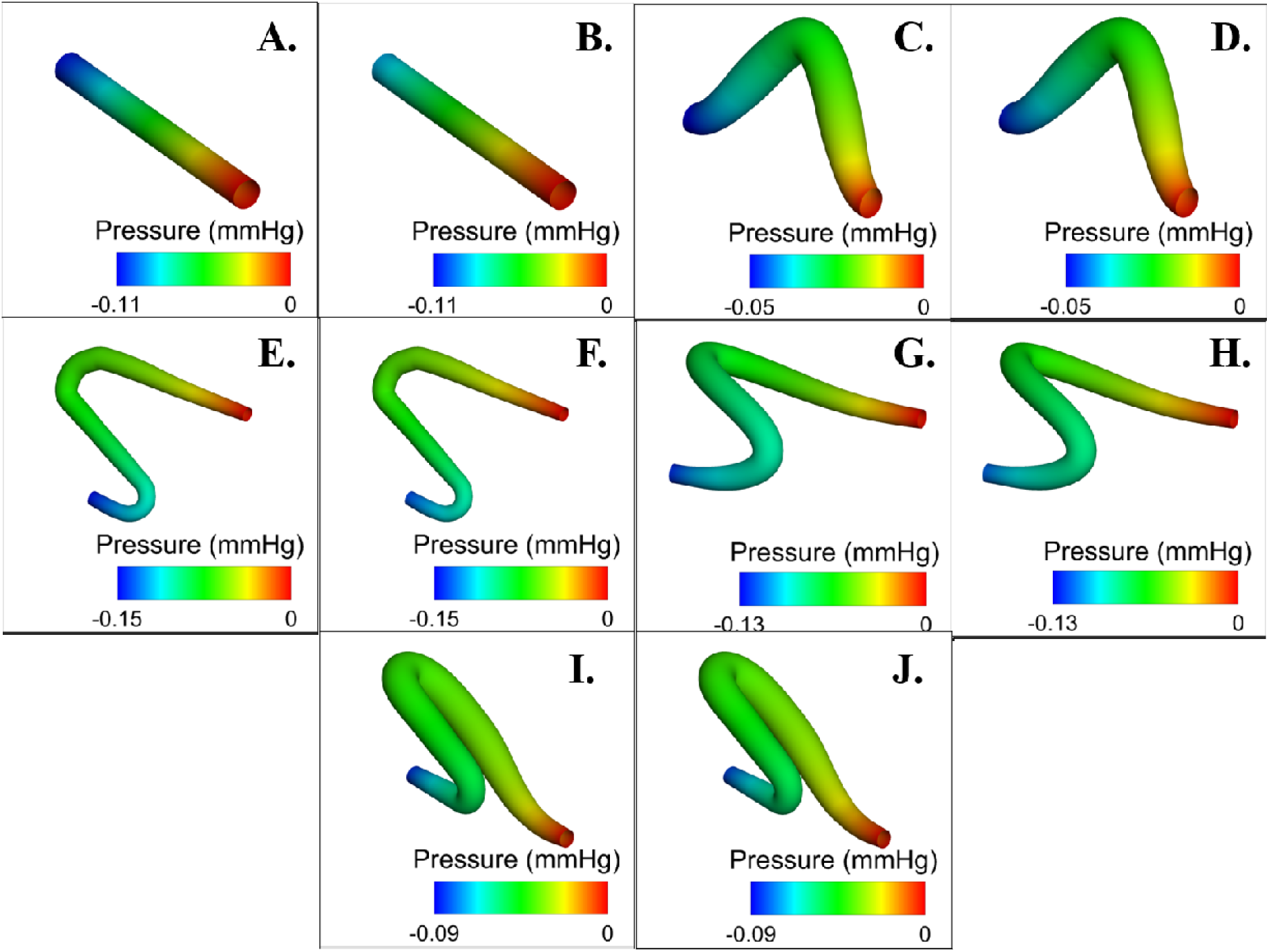
Pressure contours at peak reverse diastolic flow (t = 0.80s). Newtonian blood models: (A) Stage 1 (C) Stage 2 (E) Stage 3 (G) Stage 4 (I) Stage 5. NonNewtonian blood models: (B) Stage 1 (D) Stage 2 (F) Stage 3 (H) Stage 4 (J) Stage 5. For all geometries, the inlet diameters are 0.500 mm.

**Table 2:** Maximun peak systolic flow.

| Peak Systolic Flow |  |  |
| --- | --- | --- |
| Stage | Newtonian Pressure (mmHg) | Non-Newtonian Pressure (mmHg) |
| 1 | 0.46 | 0.27 |
| 2 | 0.24 | 0.15 |
| 3 | 0.63 | 0.41 |
| 4 | 0.56 | 0.36 |
| 5 | 0.37 | 0.25 |

Similar trends in pressure are predicted at peak reverse diastolic flow for all cases (Figure 5). Pressure values for diastolic flow are summarized in table 3. Newtonian simulations predicted larger pressure values compared to non-Newtonian simulations; however, differences in pressure between Newtonian and non-Newtonian simulations were smaller at peak reverse diastolic flow than peak systolic flow. This is likely resultant from higher viscosity values at the lower diastolic shear rates.

**Table 3:** Maximum pressures at peak reverse diastolic flow.

| Reverse Diastolic Flow |  |  |
| --- | --- | --- |
| Stage | Newtonian Pressure (mmHg) | Non-Newtonian Pressure (mmHg) |
| 1 | 0.108 | 0.090 |
| 2 | 0.055 | 0.051 |
| 3 | 0.148 | 0.139 |
| 4 | 0.131 | 0.121 |
| 5 | 0.087 | 0.085 |

### Wall shear stress comparison

Wall shear stress contour plots were generated for all geometries at peak systolic flow and peak reverse diastolic flow (Figure 6 & Figure 7). For all curved geometries during peak systolic flow (Stages 2-5), local maxima of wall shear stress occurred at the inlet, outlet, and innermost bends (Figure 6 C-J). High shear stresses at the inlet and outlet are resultant from high shear rates where luminal diameters are smallest. Additionally, local maxima in wall shear stress at the innermost bends are from large velocity values shifting from the center to closer to wall. This resulted in larger shear rates near the innermost bend (Figure 8). For all stages, Newtonian simulations predicted higher shear stresses than non-Newtonian simulations for all geometries. This produced errors larger than 40% between Newtonian and non-Newtonian simulations. This overprediction is likely explained by examining shear rates and viscosity values (Figure 3 and Figure 8). Velocity profiles for non-Newtonian cases had higher shear rates near the walls than velocity profiles for Newtonian cases. Higher shear rates resulted in decreased viscosity values near the wall for non-Newtonian cases and produced lower shear stresses than Newtonian cases. Maximum shear stresses predicted by simulations at peak systolic flow are summarized in table 4.

**Figure 6:**
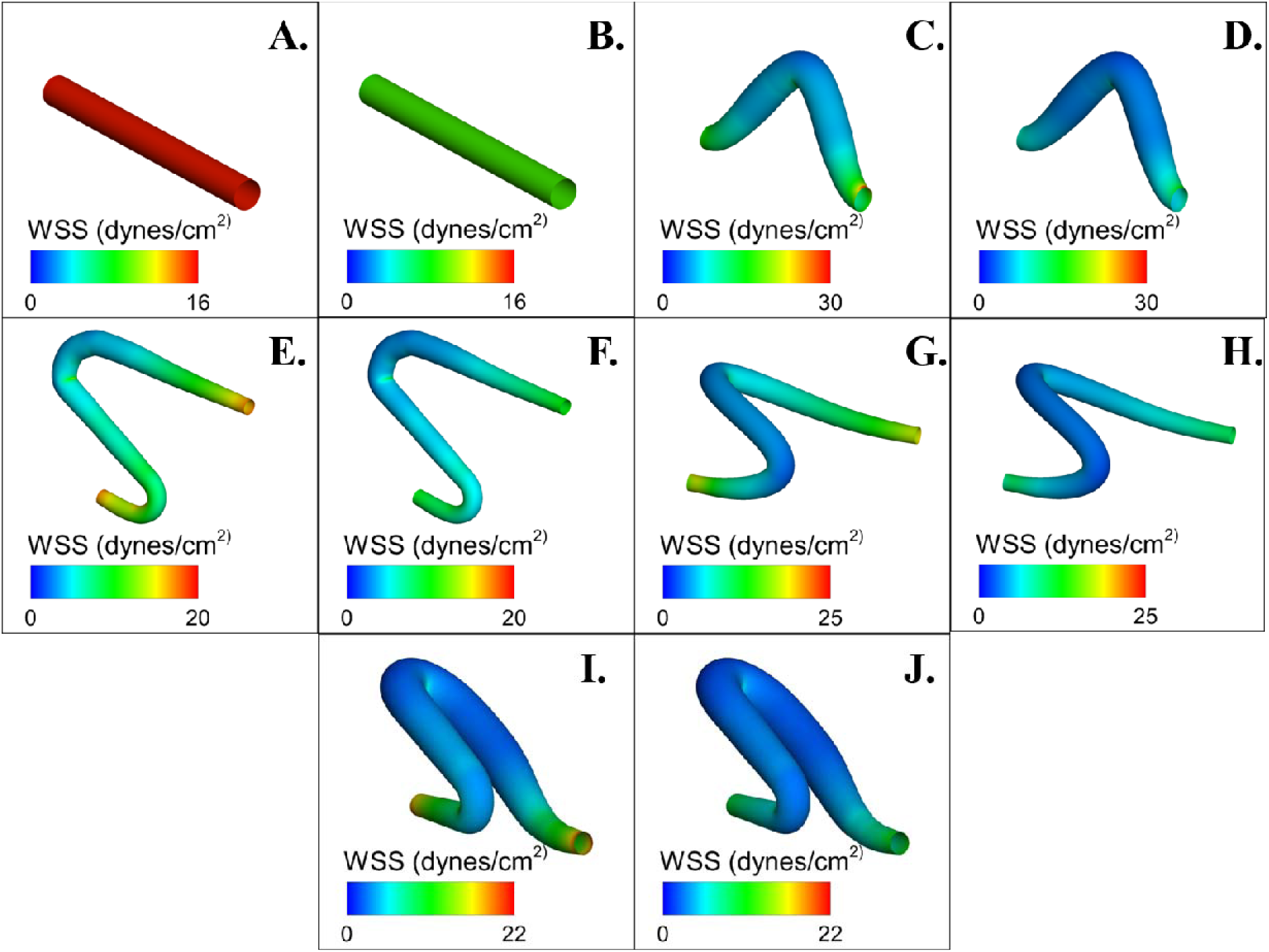
Wall shear stress contours at peak reverse diastolic flow (t = 0.80s). Newtonian models: (A) Stage 1 (C) Stage 2 (E) Stage 3 (G) Stage 4 (I) Stage 5. NonNewtonian models: (B) Stage 1 (D) Stage 2 (F) Stage 3 (H) Stage 4 (J) Stage 5. For all geometries, the inlet diameters are 0.500 mm.

**Figure 7:**
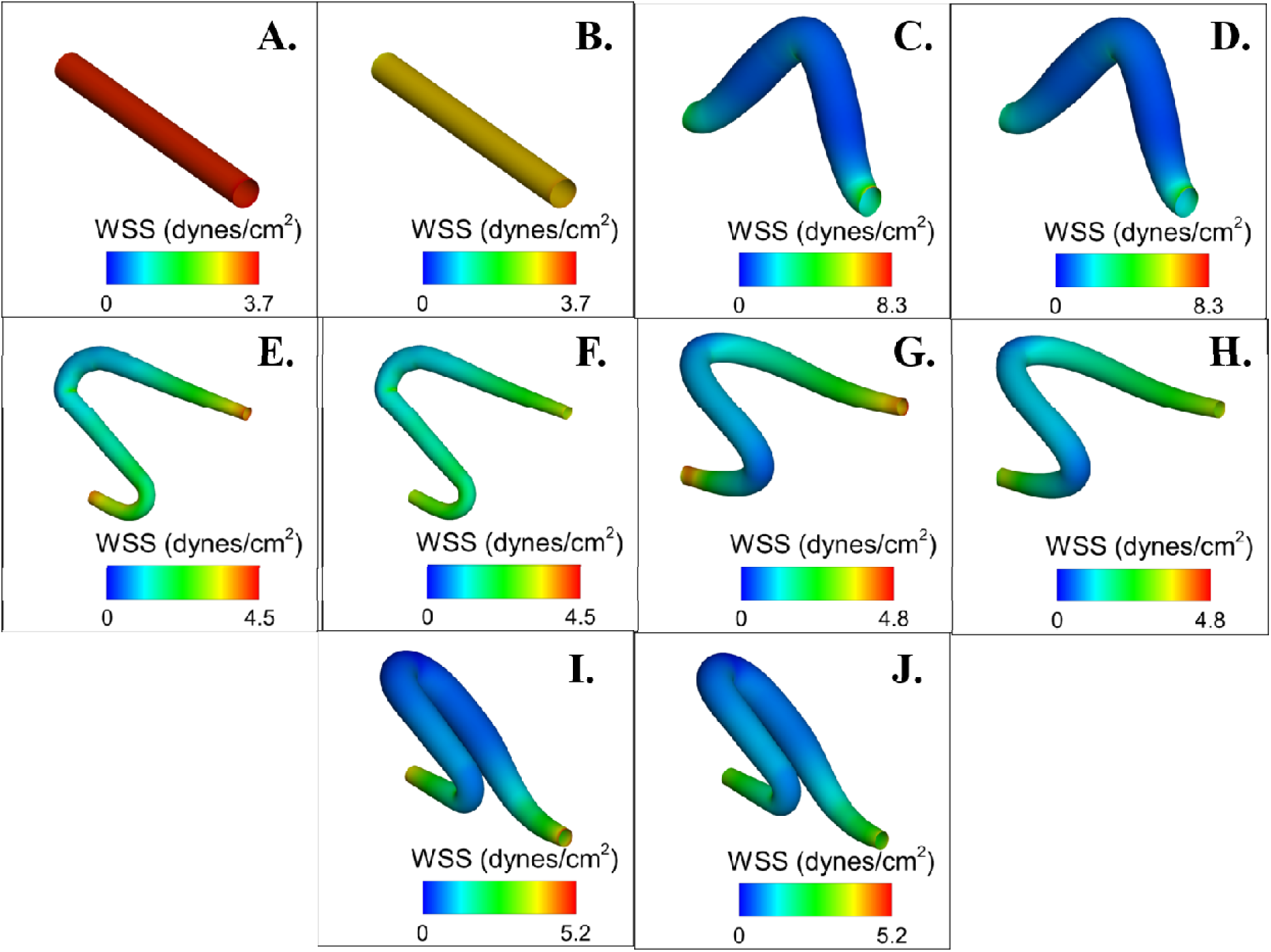
Wall shear stress contours at peak reverse diastolic flow (t = 0.80s). Newtonian blood models: (A) Stage 1 (C) Stage 2 (E) Stage 3 (G) Stage 4 (I) Stage 5. Non-Newtonian blood models: (B) Stage 1 (D) Stage 2 (F) Stage 3 (H) Stage 4 (J) Stage 5. For all geometries, the inlet diameters are 0.500 mm.

**Figure 8:**
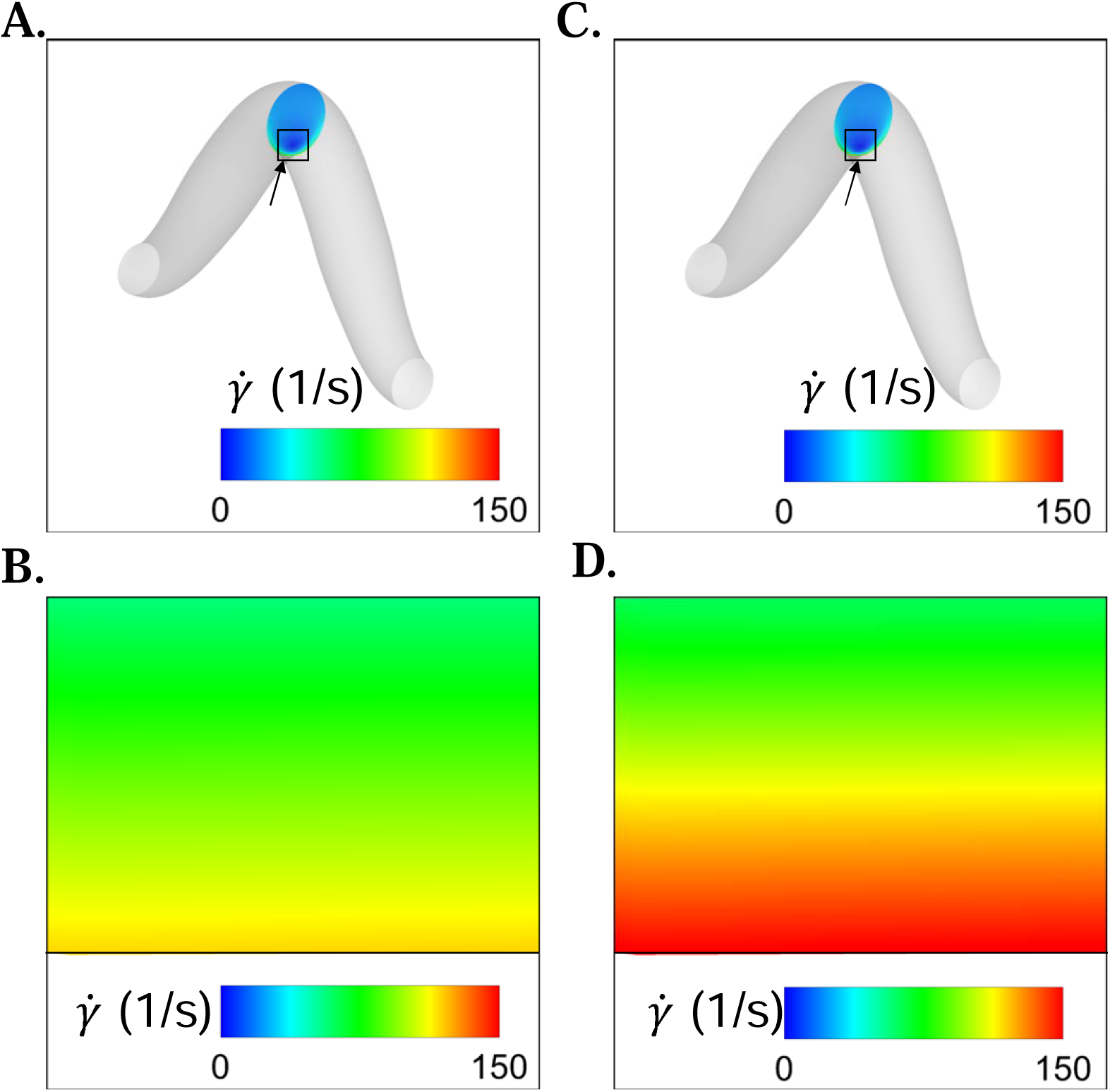
Contours of shear rate at a cross section through the bend in stage 2 geometries for Newtonian (A-B) and non-Newtonian (C-D) simulations at peak systolic flow. Black arrows in figures (A) and (C) denote the location zoomed sectionsns of shear rate near the innermost walls in Newtonian (B) and nonNewtonian (D) cases.

**Table 4:** Maximum wall shear stress at peak systolic flow.

| Peak Systolic Flow |  |  |
| --- | --- | --- |
| Stage | Newtonian Wall Shear Stress (dynes/cm<sup>2</sup>) | Non-Newtonian Wall Shear Stress (dynes/cm<sup>2</sup>) |
| 1 | 15.9 | 9.1 |
| 2 | 29.5 | 15.9 |
| 3 | 19.4 | 11.8 |
| 4 | 24.6 | 12.7 |
| 5 | 21.5 | 12.1 |

Our simulations predicted similar trends in shear stresses during peak reverse diastolic flow. For all stages, Newtonian simulations predicted larger shear stresses than non-Newtonian simulations, resulting in errors larger than 10%. This decrease in error between Newtonian and non-Newtonian flows at is due to lower shear rates caused by lower velocities. For non-Newtonian cases at low shear rates, viscosity values increase also increasing shear stresses. This resulted in a smaller difference between Newtonian and non-Newtonian shear stresses. Maximum shear stresses predicted by simulations and peak diastolic flow are summarized in table 5.

**Table 5:**
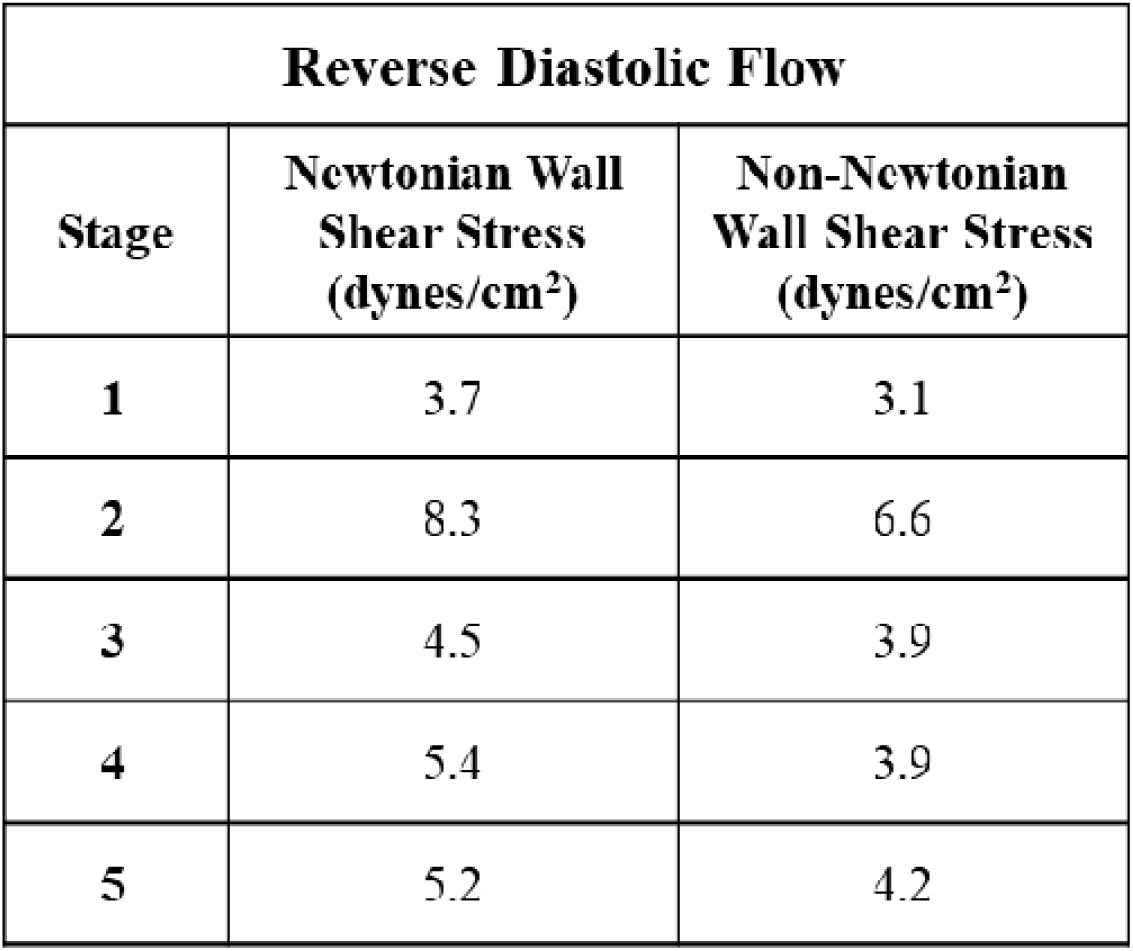
Maximum wall shear stresses at peak reverse diastolic flow.

| Reverse Diastolic Flow |  |  |
| --- | --- | --- |
| Stage | Newtonian Wall Shear Stress (dynes/cm<sup>2</sup>) | Non-Newtonian Wall Shear Stress (dynes/cm<sup>2</sup>) |
| 1 | 3.7 | 3.1 |
| 2 | 8.3 | 6.6 |
| 3 | 4.5 | 3.9 |
| 4 | 5.4 | 3.9 |
| 5 | 5.2 | 4.2 |

## DISCUSSION

Mechanical stresses are associated with the onset of blood flow within the developing embryonic heart at around three weeks in humans [20, 28]. Cells residing within the embryonic heart are sensitive to these mechanical cues, and cardiac cells respond to maintain homeostasis and progress cardiac morphogenesis [26, 29–34]. Determining physiological values of these mechanical stresses are necessary to understand cellular behavior during embryonic heart development [11, 13, 20, 35]; however, pathological values can lead to improper diagnostics, treatments, and understanding of congenital malformations that arise during heart development [36]. We present a computational model for pulsatile blood-flow through embryonic cardiac looping, accounting for the shear-thinning viscosity of blood. For all cases, we assumed a fully developed paraboloid velocity profile at the inlet. We compared results to cases where blood was treated as a Newtonian fluid with a constant viscosity, and magnitudes of wall shear stresses and pressures predicted by our models are the same order of magnitudes observed by other studies investigated embryonic cardiac flow [13, 20]. Our simulations predicted local maxima in wall shear stresses at bends in the looping geometries, like where primitive cardiac cushions develop [20], and evenly distributed wall shear stresses predicted by non-Newtonian models could affect ventricular trabeculation [37]. Additionally, our simulations discovered that Newtonian simulations using a viscosity of 0.004 Pa·s overpredicted pressures and wall-shear stresses when compared to non-Newtonian cases.

The present models assume rigid, stationary walls and therefore isolate fluid borne stresses acting on the endocardium during cardiac looping. In vivo, the embryonic heart tube is compliant and undergoes cyclic deformation; however, prior computational and experimental studies suggest that while wall motion modulates the magnitude of wall shear stress and pressure, the spatial localization of shear stress maxima—particularly at bends, inlets, and outlets—is preserved [20, 26]. By focusing on relative differences between Newtonian and non-Newtonian rheology under identical geometric and flow conditions, our approach provides mechanistic insight into how blood rheology alone biases predicted hemodynamic loads during early cardiac morphogenesis.

Our simulations predicted wall shear stresses similar to stresses observed experimentally in zebrafish and chicken embryo models of cardiac morphogenesis [38, 39], and both simulations predicted peak systolic stresses at bends in the tube close to shear stresses observed during valve development in the embryonic S-staged zebrafish heart [38]. Local maxima in shear stresses occurred at bend locations, where curvature is increased, or small luminal diameters. These locations are close to the locations of endocardial cushions, where valve development occurs later in heart morphogenesis [3, 40]. Additionally, local maxima in wall shear stress at the inlet and outlet provide insight into the development of atria and the outflow tract. For example, work demonstrated that arteries can adjust their lumen diameters to maintain shear stresses near a set optimal point [13]. Understanding what the optimal shear stress load is for normal outflow tract development could help identify conditions yielding outflow tract pathologies.

Healthy development of embryonic cardiac features are highly dependent on shear stress, and other studies have shown that cellular response to shear stress have been associated with ventricular trabeculation [12, 16], cardiac cushion formation [20], and valve differentiation [41, 42]. Gene-expression patterns associated with wall shear stress have been linked to both physiological and pathological heart formation. For example, overexpression of neuregulin-1 leads to over proliferative cardiomyocytes and can lead to hyper-trabeculation and CHDs [43]. Several other signaling pathways, such as *NOTCH, Nkx-2.5,* and *ENT1* linked to distributions and magnitudes of wall shear stress [44–46]. To this extent, non-Newtonian simulations predicted more evenly distributed wall shear stress distributions than Newtonian simulations, which could affect ventricular trabeculation. Inaccurate reported wall shear stresses hamper our understanding of physiological and pathological embryonic cardiac development. By experimentally examining genetic expression under Newtonian and non-Newtonian shear stress conditions, scientists could elucidate potential mechanisms for the onset and progression of CHDs.

In addition to wall shear stresses, cardiac pressures build during embryonic cardiac development, with rapid pressure growth after the development of cardiac valves. For example, systolic pressure drop is around 1 mmHg during the fully looped embryonic chick heart, and the pressure drop in the mature adult chicken heart is 120 mmHg.[35]. Our models do not account for valves, which form later in cardiac development, but still predict similar values to the looped embryonic heart. Pressure produces stresses in the heart wall, which influence critical signaling cascades for cardiac development and remodeling [47–50]. Newtonian models misrepresent pressure loads compared to non-Newtonian models. Pressures remain within the same order of magnitude between Newtonian and non-Newtonian simulations, but non-Newtonian models are a better representation of blood viscosity *in vivo*. Increased, and possibly pathological, ballooning is induced by high pressure loads on chamber walls. Furthermore, recent work by Dietrich *et al.* demonstrated endothelial cell proliferation and cell migration induced by flow contributed to chamber growth and ballooning in the embryonic zebrafish heart [51]. The inclusion of endocardial cushions, decreasing the luminal diameter where the tube bends, could elucidate pressure differences in pre-chamber sections of the looping heart. Our data suggest that differences in pressure values predicted by Newtonian and non-Newtonian models may induce inaccuracies in reported pressure loads during embryonic cardiac development and CHD development.

Differences in Newtonian and non-Newtonian wall shear stresses and pressures are likely resultant of differences in velocity profiles between Newtonian and non-Newtonian cases and decreased viscosities at higher shear rates in non-Newtonian cases. Experimental models utilizing flow imaging techniques often use a Newtonian viscosity approximation to scale calculated shear rates to shear stresses and to evaluate pressures [39]. Viscosities ranging from 0.001 Pa·s to 0.008 Pa·s have been used, resulting in a wide range of stresses reported [38]. Circumferential strains induced by luminal pressure loads and wall shear stresses affect gene expression in both endothelial cells and cardiomyocytes [52]. Using a non-Newtonian model to predict stresses and pressures is more representative of the behavior of embryonic blood during embryonic cardiac development. Stresses and pressures predicted by non-Newtonian models could improve *in vitro* tissue engineered models of embryonic cardiac development and could better evaluate mechanotransducive sensitive pathways in cardiac cells. Furthermore, inaccurate magnitudes of wall shear stress and pressures from Newtonian models could be closer to pathological values and could misevaluate flow conditions during CHD formation and progression.

We conclude that numerical simulations of blood flow through stages of cardiac looping using a Newtonian model of blood viscosity overpredict wall shear stresses and pressures compared to a non-Newtonian model of blood viscosity. Simulations predicted local maxima in wall shear stresses where atria, outflow tract, and cardiac cushions develop during later stages of heart looping. Additionally, non-Newtonian simulations more evenly distributed wall shear stresses, which could affect ventricular trabeculation. Predicted upstream pressures could modulate ventricular ballooning. Future experimental studies should examine the relationship between non⍰Newtonian flow conditions and downstream mechanobiological responses in cardiac cells. We suggest that non⍰Newtonian viscosity models be used alongside Newtonian approximations when evaluating embryonic hemodynamics, as they provide a more physiologically relevant representation of blood behavior during cardiac morphogenesis. We suggest that non-Newtonian viscosity models of embryonic blood be used in alongside Newtonian approximations when evaluating hemodynamics during embryonic cardiac development, and that non-Newtonian models can aid in our knowledge of flow patterns and stresses during physiological and pathological cardiac morphogenesis.

## ETHICS, CONSENT TO PARTICIPATE, AND CONSENT TO PUBLISH DECLARATIONS

Not Applicable.

